# Neurodevelopmental processes in the prefrontal cortex derailed by chronic HIV-1 viral protein exposure

**DOI:** 10.1101/2021.09.02.458765

**Authors:** Kristen A. McLaurin, Hailong Li, Rosemarie M. Booze, Charles F. Mactutus

## Abstract

Due to the widespread access to, and implementation of, combination antiretroviral therapy, individuals perinatally infected with human immunodeficiency virus type 1 (HIV-1) are living into adolescence and adulthood. Perinatally infected adolescents living with HIV-1 (pALHIV) are plagued by progressive, chronic neurocognitive impairments; the pathophysiological mechanisms underlying these deficits, however, remains understudied. A longitudinal experimental design from postnatal day (PD) 30 to PD 180 was utilized to establish the development of pyramidal neurons, and associated dendritic spines, from layers II-III of the medial prefrontal cortex (mPFC). Three putative neuroinflammatory markers (i.e., IL-1β, IL-6, and TNF-α) were evaluated early in development (i.e., PD 30) as a potential mechanism underlying synaptic dysfunction in the mPFC. Constitutive expression of HIV-1 viral proteins induced prominent neurodevelopmental alterations, independent of biological sex, in pyramidal neurons from layers II-III of the mPFC. Specifically, HIV-1 transgenic rats exhibited prominent deficits in dendritic and synaptic pruning, a developmental decrease in synaptic connectivity, and an age-related decline in synaptic efficacy. Examination of dendritic spine morphology revealed an age-related population shift towards a more immature dendritic spine phenotype in HIV-1 transgenic animals. There was no compelling evidence for neuroinflammation in the mPFC during early development. Understanding the neural mechanisms underlying chronic neurocognitive impairments in pALHIV may afford a key target for innovative therapeutics and cure strategies; an urgent need given the growing population of pALHIV.

## 1. Introduction

Reports of children diagnosed with acquired immune deficiency syndrome (AIDS) occurred early in the pandemic [1-3]. Invariably, the disease was associated with high mortality rates [4-5]. However, with widespread access to, and implementation of, combination antiretroviral therapy (cART), pediatric human immunodeficiency virus type 1 (HIV-1) evolved from a fatal disease to a chronic disease [e.g., 6-7]. Consequently, the epidemiological features of pediatric HIV-1 have shifted, resulting in a growing population of perinatally infected adolescents living with HIV-1 (pALHIV). Indeed, approximately 2.78 million children and adolescents (0-19 years of age) are living with either perinatally or horizontally acquired HIV-1 [8]. Despite the disproportionate number of adolescents affected by HIV-1 worldwide, neurodevelopmental outcomes of pALHIV remain understudied [9].

Neurodevelopment, a dynamic, albeit protracted, process beginning during the gestational period and continuing through late adolescence, refers to the maturation of neural circuits that influence cognitive functioning [for review, 10]. Early pre- and postnatal brain development occurs via a precisely orchestrated and timed series of progressive events, including neuronal production, migration, and differentiation, as well as myelination and synaptogenesis [10-11]; events which permit the establishment of neural circuits. Subsequent regressive events, including programmed cell death [12-14], synaptic pruning [15-16], and dendritic pruning [17], aid in the refinement of neural circuits resulting in the formation of mature connections. Notably, the time course of such progressive and regressive events varies considerably across the cerebral cortex, whereby the prefrontal cortex (PFC) is one of the latest.

Whereas the basic cytoarchitecture of the PFC is defined by birth in humans [for review, 18] and postnatal day (PD) 10 in rodents [19], its fine development continues through adolescence and young adulthood. Specifically, following neuronal proliferation and migration, cortical nerve cells extend their axons, undergo dendritic arborization, and form dendritic spines [20-21], enabling the assembly of immature neuronal circuits. The initial patterning of neural circuits in the PFC reflects exuberant connectivity, evidenced by the formation of an excessive number of synaptic contacts; synaptic contacts which are subsequently pruned during adolescence and young adulthood [15-16; 22], coinciding with the development of higher-order cognitive processes. Chronic HIV-1 viral protein exposure may disrupt these events, fundamentally altering neurodevelopment in the PFC resulting in profound progressive neurocognitive impairments [23-24].

Indeed, chronic neurocognitive impairments [25] are a highly prevalent [26] deleterious consequence of perinatally acquired HIV-1. Cross-sectional studies have provided a wealth of knowledge on chronic neurocognitive impairments, characterized by prominent deficits in cognitive functions that are dependent upon the integrity of the PFC [e.g., Processing Speed: 27-28; Attention: 29; Executive Function: 30]. Evaluating neurodevelopmental outcomes, however, necessitates the utilization of a longitudinal experimental design; extrapolating cross-sectional findings to developmental processes is inferentially fraught [31]. Thus, repeated evaluation of neurocognitive function across time has been essential for establishing the progressive nature of chronic neurocognitive impairments associated with perinatally acquired HIV-1 [23-24; 32]. Longitudinal experimental designs, however, remain underutilized for the investigation of the pathophysiological mechanisms underlying progressive chronic neurocognitive impairments, albeit with the exception of a recent manuscript [33].

To address this knowledge gap, two interrelated aims were investigated. First, a longitudinal experimental design (PD 30 to PD 180) was utilized to establish the development of pyramidal neurons, and associated dendritic spines, from layers II-III of the medial PFC (mPFC). Profound synaptic dysfunction has been observed in HIV-1 seropositive individuals [34-35] and biological systems used to model key aspects of HIV-1 [e.g., 24; 36-37]; neurodevelopmental alterations in pyramidal neurons and/or dendritic spines, however, have not been systematically evaluated. Given the fundamental role of experience on neuronal and dendritic spine morphology [e.g., 38-40], synaptic dysfunction was evaluated in HIV-1 Tg and control animals with no prior cognitive or behavioral testing. Second, neuroinflammatory markers (i.e., IL-1β, IL-6, and TNF-α) were evaluated in the PFC at PD 30 to evaluate a potential mechanism underlying synaptic dysfunction [e.g., IL-6: 41]. Understanding the neural mechanisms underlying chronic neurocognitive impairments in pALHIV may afford a key target for innovative therapeutics and cure strategies; an urgent need given the growing population of pALHIV.

## 2. Materials and Methods

### 2.1. Animals

Control (Fischer F344/N; Envigo Laboratories Inc., Indianapolis, IN) animals were delivered to the animal vivarium and pair- or group-housed with animals of the same sex until sacrifice. HIV-1 Tg animals, originally reported by Reid et al [42], were bred on a Fischer F344/N background at the University of South Carolina by pairing a control female with a HIV-1 Tg male. At approximately PD 24, HIV-1 Tg animals were weaned and pair- or group-housed with animals of the same sex until sacrifice. To preclude violation of the independence of observations assumption, the goal was to sacrifice no more than two rats (i.e., one male and one female) from each litter for an individual time point.

AAALAC-accredited facilities and guidelines established in the Guide for the Care and Use of Laboratory Animals of the National Institutes of Health were utilized for the maintenance of HIV-1 Tg and control animals. Animal vivarium environmental conditions were targeted at: 21°± 2°C, 50% ± 10% relative humidity and a 12-h light:12-h dark cycle with lights on at 0700 h (EST). The Institutional Animal Care and Use Committee (IACUC) at the University of South Carolina approved the project protocol (Federal Assurance, #D16-00028).

### 2.2. Experiment #1: Neurodevelopmental Alterations in the Medial Prefrontal Cortex (mPFC)

Pyramidal neurons, and associated dendritic spines, from layers II-III of the mPFC were examined to establish neurodevelopmental alterations in the mPFC. HIV-1 Tg and control animals were sacrificed every 30 days from PD 30 to PD 180 utilizing samples sizes of n=40 for each age (i.e., Control: Male, n=10, Female, n=10; HIV-1 Tg: Male, n=10, Female, n=10). Ages were selected based on deficits observed in previous imaging [33] and cognitive [23; 43-44] studies. Specifically, the earliest timepoint (i.e., PD 30) was selected following reports that presence of the HIV-1 transgene induced alterations in brain volume development early in life (i.e., 5-9 weeks of age [33]). The study endpoint (i.e., PD 180) reflects an age whereby the HIV-1 Tg rat exhibits prominent neurocognitive impairments relative to control animals [e.g., 23; 43-44].

#### 2.2.1. Body Weight

Body weight, a measurement of somatic growth, was assessed immediately prior to sacrifice. Independent of biological sex, HIV-1 Tg animals weighed significantly less than control animals from PD 30 to PD 180.

Body weight for both male and female animals, independent of genotype, was well-described by a one-phase association (Supplementary Figure 1; R^2^s≥0.86). For male animals, presence of the HIV-1 transgene significantly altered the plateau of the function (*F*(1,106)=14.1, *p*≤0.001), but not the rate of growth (i.e., no statistically significant effect of genotype on the rate constant, K (*p*>0.05)). For female animals, HIV-1 Tg animals exhibited a significantly lower y-intercept (*F*(1,108)=10.4, *p*≤0.001) and plateau (*F*(1,108)=7.2, *p*≤0.01), but no statistically significant differences in the rate of growth (*p*>0.05). At sacrifice, therefore, there was strong evidence for the relative health of the HIV-1 Tg rat.

#### 2.2.2. Estrous Cycle Tracking

A vaginal lavage was conducted immediately prior to sacrifice to evaluate estrous cyclicity. Cellular cytology was examined in vaginal smears, as previously described [45], under a 10x light microscope. The predominant cell type was utilized to determine the estrous cycle stage [46-47], whereby the diestrus phase was characterized by the predominance of leukocytes [48]. The goal was to sacrifice female animals during the diestrus phase of the estrous cycle to reduce potential variability due to hormonal cycle.

#### 2.2.3. Ballistic Labeling Technique

Methodology for the ballistic labeling technique was initially described by Seabold et al. [49] and was adapted for use in our laboratory by Roscoe et al. [37]; a protocol that was further refined by Li et al. [50].

In brief, the DiI/Tungsten bead tubing was created by dissolving polyvinylpyrrolidone (PVP; 100 mg) in ddH_2_O (10 mL). Tefzel tubing (Bio-Rad, Hercules, CA, USA) was filled with the PVP solution for 20 minutes and then expelled. DiOlistic cartridges were prepared by dissolving tungsten beads (170 mg; Bio-Rad) with 99.5% methylene chloride (250 μL; Sigma-Aldrich, St. Louis, MO, USA); the tungsten bead suspension was vortexed to mix thoroughly. Additionally, DiIC18(3) dye (6 mg; Invitrogen, Carlsbad, CA, USA) was dissolved in 99.5% methylene chloride (300 μL) and vortexed. The tungsten bead suspension was pipetted onto a glass slide and allowed to dry. After pipetting the DiIC18(3) dye solution on top of the tungsten bead suspension, the two layers were mixed; the tungsten bead suspension/DiIC18(3) dye mixture were split into two 1.5 mL centrifuge tubes filled with ddH_2_O and sonicated. The homogenized mixture in the 1.5 mL centrifuge tubes was combined into a 15 mL conical tube, further sonicated, drawn into the PVP-coated Tefzel tubing, and fed into the tubing preparation station (Bio-Rad). After the PVP-coated Tefzel tubing with the homogenous tungsten bead suspension/DiIC18(3) dye mixture (i.e., DiOlistic cartridge) rotated for one minute, all water was removed. Rotating continued for an additional 30 minutes under nitrogen gas (0.5 liters per minute). Once dried, the DiOlistic cartridges were cut into 13 mm lengths and stored in the dark until subsequent use.

HIV-1 Tg and control animals were deeply anesthetized using 5% sevoflurane (Abbot Laboratories, North Chicago, IL, USA) and transcardially perfused. The rat brain was removed, postfixed in 4% paraformaldehyde for 10 minutes and cut coronally (500 μm) using a rat brain matrix (ASI Instruments, Warren, MI, USA). Coronal slices were placed into a 24 well-plate with 100 mM phosphate-buffered saline (PBS; 1 mM). Before ballistic labeling, the PBS was removed from the well-plate.

Ballistic labeling was conducted using the Helios gene gun (Bio-Rad), which was loaded with previously prepared DiOlistic cartridges. A piece of filter paper was placed between the two mesh screens on the Helios gene gun. For ballistic delivery, helium gas flow was adjusted to 90 pounds per square inch and the applicator was placed approximately 2.5 cm away from the brain slices. After ballistic delivery, slices were washed three times in 100 mM PBS and stored in the dark at 4°C for three hours. Brain slices were transferred onto a glass slide, mounted using Pro-Long Gold Antifade (Invitrogen), and cover slipped (#1 cover slip; Thermo Fisher Scientific, Waltham, MA, USA). Slides were stored in the dark at 4°C until imaging.

#### 2.2.4. Confocal Imaging of Pyramidal Neurons

Pyramidal neurons from layers II-III of the mPFC, located approximately 3.7 mm to 2.2 mm anterior to Bregma [51], were imaged for analysis (Figure 1). Z-stack images (60x oil objective (n.a.=1.4) with Z-plane interval of 0.15 µm) of three pyramidal neurons from each animal were obtained using a Nikon TE-200E confocal microscope and Nikon’s EZ-C1 software (version 3.81b). The DiI flurophore was excited using a green helium-neon laser with an emission of 543 nm.

**Figure 1.**
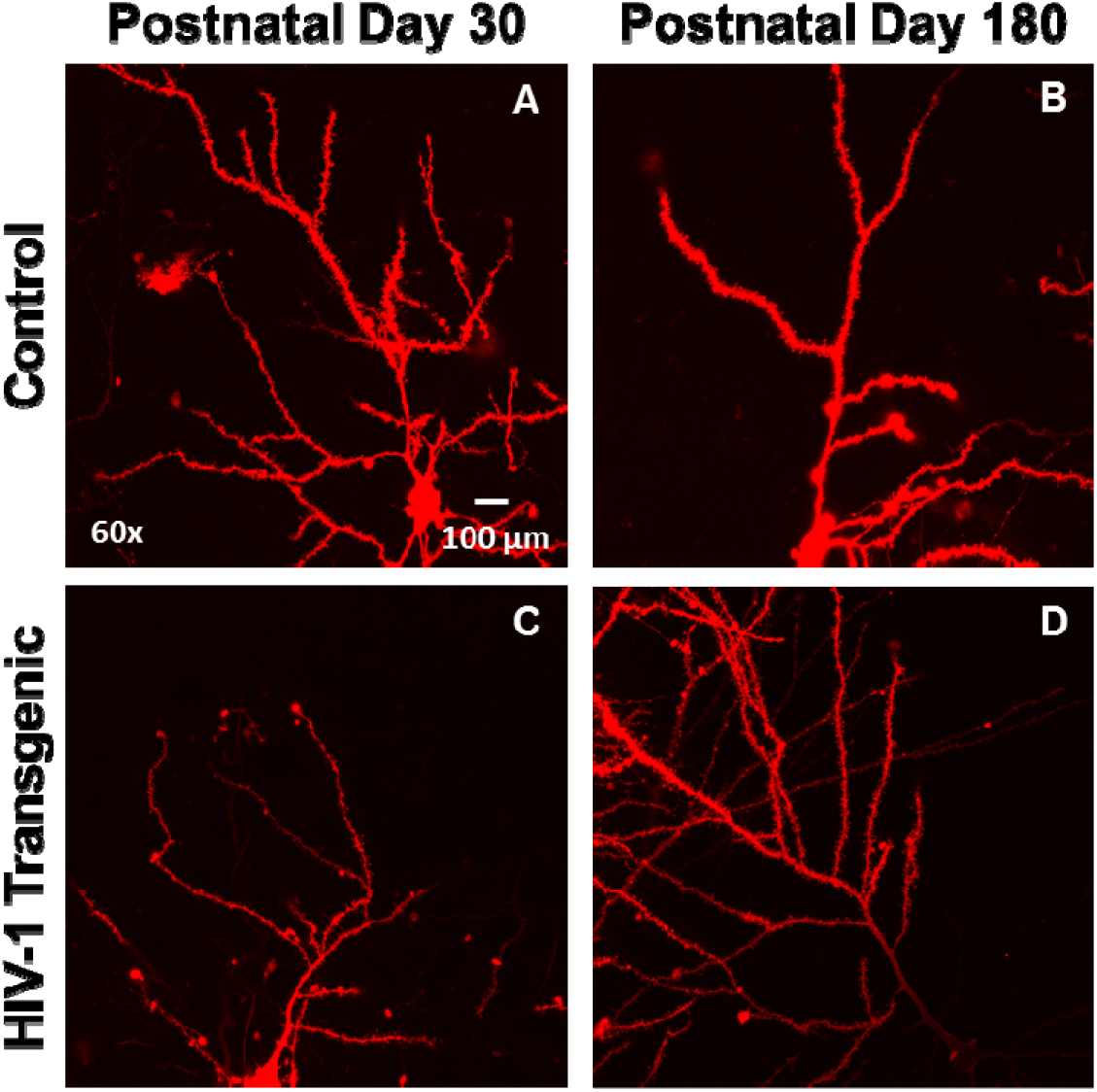
Representative images of DiOlistically labeled pyramidal neurons in layers II-III of the medial prefrontal cortex (mPFC) in control (**A-B**) and HIV-1 transgenic (Tg; **C-D**) animals at postnatal day (PD) 30 (**A**,**C**) and PD 180 (**B**,**D**).

#### 2.2.5. Neuronal Analysis and Spine Quantification

Neuronal morphology and dendritic spines were analyzed using sophisticated neuronal reconstruction software (Neurolucida 360, MicroBrightfield, Williston, VT). Selection criteria (e.g., continuous dendritic staining, low background/dye clusters), illustrated in Li et al. [50], were used to select one neuron from each animal for analysis. Neurons not meeting the selection criteria were not included in the analysis, yielding the following sample sizes: Control: PD 30, *n=*19 (male, *n*=9, female, *n*=10), PD 60, *n*=19 (male, *n*=10, female, *n*=9), PD 90, *n*=18 (male, *n*=10, female, *n*=8), PD 120, *n*=17 (male, *n*=9, female, *n*=8), PD 150, *n*=18 (male, *n*=8, female, *n*=10), PD 180, *n*=20 (male, *n*=9, female, *n*=11); HIV-1 Tg: PD 30, *n*=19 (male, *n*=10, female, *n*=9), PD 60, *n*=19 (male, *n*=10, female, *n*=9), PD 90, *n*=18 (male, *n*=8, female, *n*=10), PD 120, *n=*20 (male, *n*=10, female, *n*=10), PD 150, *n*=19 (male, *n*=9, female, *n*=10), PD 180, *n*=18 (male, *n*=9, female, *n*=9). Following selection, the apical dendrite of pyramidal neurons from layers II-III of the mPFC was examined.

Morphological characteristics of pyramidal neurons from layers II-III of the mPFC were examined using three approaches, including apical dendrite length, Sholl analysis [52] and centrifugal branch ordering. The classical Sholl analysis affords an opportunity to examine neuronal arbor complexity by counting the number of dendritic intersections occurring at successive concentric circles placed fixed distances (i.e., 10 µm) from the soma. A centrifugal branch ordering scheme assigned each dendrite with a branch order by counting the number of segments traversed; an assessment of dendritic branching complexity.

The morphology of dendritic spines was characterized by three parameters, including volume (µm^3^), backbone length (µm), and head diameter (µm). Boundary conditions for each parameter were established using well-accepted previously published results (volume, 0.05 to 0.85 µm: [53]; backbone length, 0.4 to 4.0 µm: [54-55]; head diameter, 0 to 1.2 µm: [56]). Dendritic spines recognized by the sophisticated neuronal software, but failing to meet any of the boundary conditions, were excluded from dendritic spine morphological analyses.

### 2.3. Experiment #2: Neuroinflammatory Markers in the Medial Prefrontal Cortex (mPFC)

Neuroinflammation was assessed in the mPFC of HIV-1 Tg (*n*=20; Male, *n*=10, Female, *n*=10) and control (*n*=20; Male, *n*=10, Female, *n*=10) animals at PD 30 using real-time polymerase chain reaction (RT-PCR) and enzyme-linked immunosorbent assays (ELISA).

#### 2.3.1. Sacrifice

HIV-1 Tg and control animals were humanely euthanized and sacrificed by rapid decapitation. Brains were removed and the frontal cortex was dissected and stored at -80°C until being utilized for either RT-PCR or ELISA.

#### 2.3.2. Real-Time Polymerase Chain Reaction (RT-PCR)

The RNeasy FFPE kit (QIAGEN, Germantown, MD) was utilized to isolate the total RNA from 30 mg of mPFC tissue. Using the Invitrogen (Carlsbad, CA) Cloned AMV First-Strand cDNA Synthesis Kit, one μg of total RNA from each sample was converted into cDNA. Details regarding the cDNA synthesis reaction mixture are available in McLaurin et al. [24].

Subsequently, three neuroinflammatory markers, including IL-1β, IL-6, and TNF-α were quantified using real-time PCR and the SsoAdvanced Universal SYBR Green Supermix Kit (BIO-RAD, Hercules, CA). β-actin was used as an internal control. Methodological details are available in McLaurin et al. [24]. Animals failing to reach threshold after 40 cycles were considered to have undetectable gene expression and accordingly these censored data were not included in the figures or statistical analysis, yielding Control, *n=*20, male, *n=*10, female, *n*=10, and HIV-1 Tg *n=*13-19, male, *n=*6-10, female, *n*=7-9, dependent upon gene. Data were analyzed using Intuitive Opticon Monitor TM software.

#### 2.3.3. Enzyme-Linked Immunosorbent Assay (ELISA)

Brain tissue lysates were prepared from 12-54 mg of fresh frozen PFC tissue. RIPA Lysis and Extraction Buffer (1x, Thermo Scientific, Waltham, MA) was added to extract protein from the brain tissue. Tissue was disrupted using hand sonication. Lysates were subsequently centrifuged (14400 rpm at 4°C for 20 minutes) and the supernatant was collected. A commercial ELISA kit was utilized to evaluate levels of IL-1β (R&D Systems, Inc., Minneapolis, MN), the cytokine with the highest levels of expression in RT-PCR. ELISA was analyzed in duplicate.

### 2.4. Statistical Analysis

Analysis of variance (ANOVA) and regression techniques ((SAS/STAT Software 9.4, SAS Institute, Inc., Cary, NC; SPSS Statistics 27, IBM Corp., Somer, NY; GraphPad Software, Inc., La Jolla, CA) were utilized for the statistical analysis of all data. Statistical significant was set at an alpha criterion of *p*≤0.05.

Multiple dependent measures were utilized to evaluate the impact of chronic HIV-1 viral protein exposure, biological sex, and/or age on regressive processes, including dendrite and synaptic pruning. To evaluate the progression of neuronal arbor complexity, area under the Sholl curve was derived from the classic Sholl intersection profile for each animal at every age. A mixed-model ANOVA with a compound symmetry covariance structure (PROC MIXED; SAS/STAT Software 9.4) was utilized to statistically analyze the area under the Sholl curve. The classic Sholl intersection profile was analyzed using a mixed-model ANOVA with a variance components covariance structure (PROC MIXED; SAS/STAT Software 9.4), as recommended by Wilson et al. [57]. The total number of dendritic branches, an index of dendritic branching complexity, and dendritic spines, a measure of excitatory synapses, were evaluated using a generalized linear mixed effects model with a Poisson distribution and unstructured covariance structure (PROC GLIMMIX; SAS/STAT Software 9.4). Complementary regression techniques (GraphPad Software, Inc.) were also used to assess the development of apical dendrite length, neuronal arbor complexity, dendritic branching complexity, and excitatory synapses.

Synaptic connectivity was assessed by evaluating the number of dendritic spines between each successive radii. Three measures of dendritic spine morphology, including dendritic spine volume, backbone length, and head diameter, were analyzed by examining the number of dendritic spines within each bin. A generalized linear mixed effects model with a Poisson distribution and an unstructured covariance pattern was conducted using PROC GLIMMIX (SAS/STAT Software 9.4) for both synaptic connectivity and dendritic spine morphological parameters. The statistical analyses for dendritic spine backbone length were conducted for bins from 0.4 μm to 2.0 μm. Furthermore, the statistical analyses for dendritic spine head diameter were conducted for dendritic spines with a head diameter greater than 0.01 μm. Radii and bin served as within-subjects factors, as appropriate, whereas genotype (HIV-1 Tg vs. Control), biological sex (Male vs. Female) and age (Synaptic Connectivity: PD 30 vs. PD 180; Dendritic Spine Morphology: PD 30, PD 60, PD 90, PD 120, PD 150, PD 180) served as between-subjects factors.

Putative neuroinflammatory markers were examined using two complementary measures, including RT-PCR and ELISA. For RT-PCR, gene expression was investigated using ΔCT, which was calculated by subtracting the gene of interest from the internal control (i.e., β-actin). ΔCT was analyzed independently for each neuroinflammatory marker using a mixed-model ANOVA (PROC MIXED; SAS/STAT Software 9.4). Interactions were further evaluated using simple effects in the context of the overall analysis. For ELISA, the concentration was averaged across the two duplicates to account for the nested data structure [58] and analyzed using a mixed-model ANOVA (SPSS Statistics 27). Genotype (HIV-1 Tg vs. Control) and biological sex (Male vs. Female) served as between-subjects factors.

## 3. Results

### 3.1. Experiment #1: Neurodevelopmental Alterations in the Medial Prefrontal Cortex (mPFC)

#### 3.1.1. Presence of HIV-1 viral proteins results in abnormal development and patterning of dendritic branches

HIV-1 Tg animals displayed prominent alterations in the development and patterning of apical dendrites in pyramidal neurons in layers II-III of the mPFC, evidenced by three complementary measures of neuronal morphology (i.e., Dendrite Length, Figure 2A; Sholl Analysis, Figure 2B; Dendritic Branching, Figure 2C).

**Figure 2.**
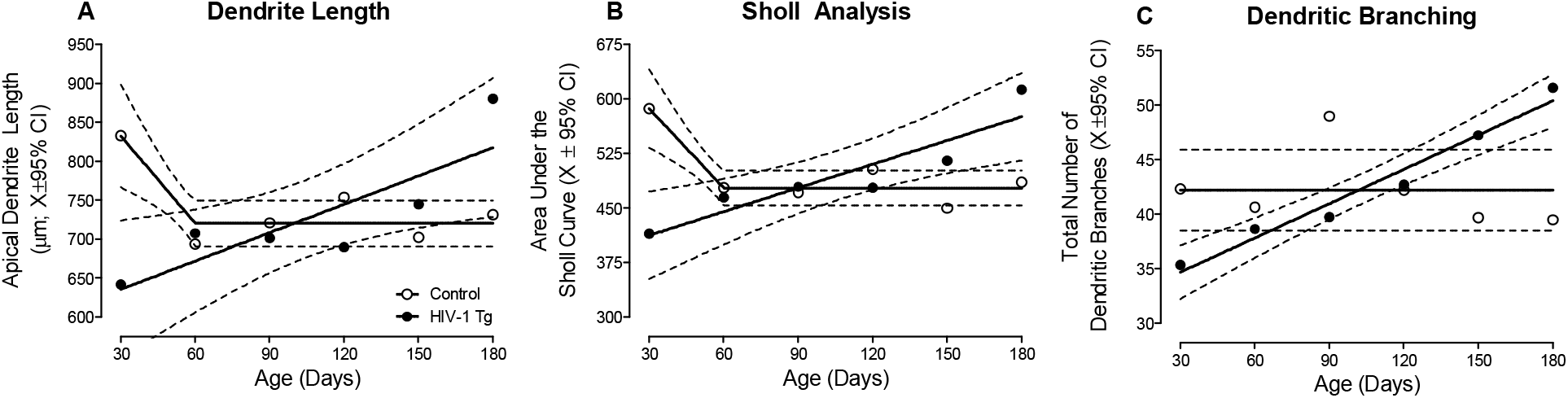
Chronic HIV-1 viral protein exposure alters neuronal development in pyramidal neurons from layers II-III of the medial prefrontal cortex, evidenced by measurements of apical dendrite length (**A**), Sholl analysis (**B**) and dendritic branching (**C**). To evaluate the progression of neuronal arbor complexity, the area under the Sholl curve was derived from the classic Sholl intersection profile. HIV-1 Tg animals exhibited a linear increase in dendrite length, the area under the Sholl curve, and the total number of dendritic branches across development; a developmental trajectory that was in sharp contrast to observations in control animals. Collectively, results support aberrant dendritic pruning in HIV-1 Tg animals relative to controls. Data are illustrated a function of genotype (Control vs. HIV-1 Tg) and age (X ± 95% confidence intervals).

A differential progression, dependent upon genotype, was observed in both apical dendrite length (Figure 2A) and the area under the Sholl curve (Figure 2B; Genotype x Age interaction, *F*(5,200)=2.6, *p*≤0.03). Specifically, in control animals, apical dendrite length and the classical Sholl analysis revealed initial neuronal arbor elaboration (i.e., PD 30) followed by dendritic pruning during adolescence (i.e., PD 60) and the stabilization of the neuronal arbor (i.e., PD 90-180; Best Fit: Segmental Linear Regression, R^2^≥0.84 and R^2^≥0.87 for apical dendrite length and area under the Sholl curve, respectively). In HIV-1 Tg animals, however, a linear increase in both apical dendrite length and neuronal arbor complexity was observed across development (Best Fit: First-Order Polynomial, R^2^≥0.71 and R^2^≥0.83 for apical dendrite length and area under the Sholl curve, respectively). Critically, with regards to the Sholl analysis, examination of the Sholl intersection profile revealed consistent observations (Supplementary Figure 2; Genotype x Age x Radius interaction, *F*(5, 5352)=3.1, *p*≤0.01).

Furthermore, profound alterations in the branching complexity of dendrites were observed in HIV-1 Tg animals relative to controls (Figure 2C; Genotype x Age interaction, *F*(5, 200)=2.4, *p*≤0.04)). HIV-1 Tg animals exhibited a progressive increase in the total number of dendritic branches throughout development (Best Fit: First-Order Polynomial, R^2^≥0.96); a sharp contrast to the stable number of total dendritic branches observed in control animals. Taken together, HIV-1 Tg animals exhibited enhanced neuronal arbor and branching complexity across development supporting aberrant dendritic pruning.

#### 3.1.2. Prominent alterations in synaptogenesis were observed in HIV-1 Tg rats

Given that dendritic spines typically contain a single type 1 asymmetric synapse [59], the total number of dendritic spines may serve as a proxy for the number of excitatory synapses [for review, 60]. A differential progression, dependent upon genotype, was observed in the total number of dendritic spines across development (Figure 3; Genotype x Age Interaction, *F*(5, 200)=9.9, *p*≤0.001). Specifically, control rats, independent of biological sex, exhibited an initial overproduction of dendritic spines (i.e., PD 30) followed by pruning during adolescence (i.e., PD 60) and the stabilization of dendritic spines during adulthood (i.e., PD 90-180; Best Fit: Segmental Linear Regression, R^2^≥0.87). In sharp contrast, HIV-1 Tg animals exhibited a linear increase in the number of dendritic spines across development (Best Fit: First-Order Polynomial, R^2^≥0.92). Thus, independent of biological sex, presence of the HIV-1 transgene alters synaptogenesis, including the establishment, maintenance, and elimination of synapses.

**Figure 3.**
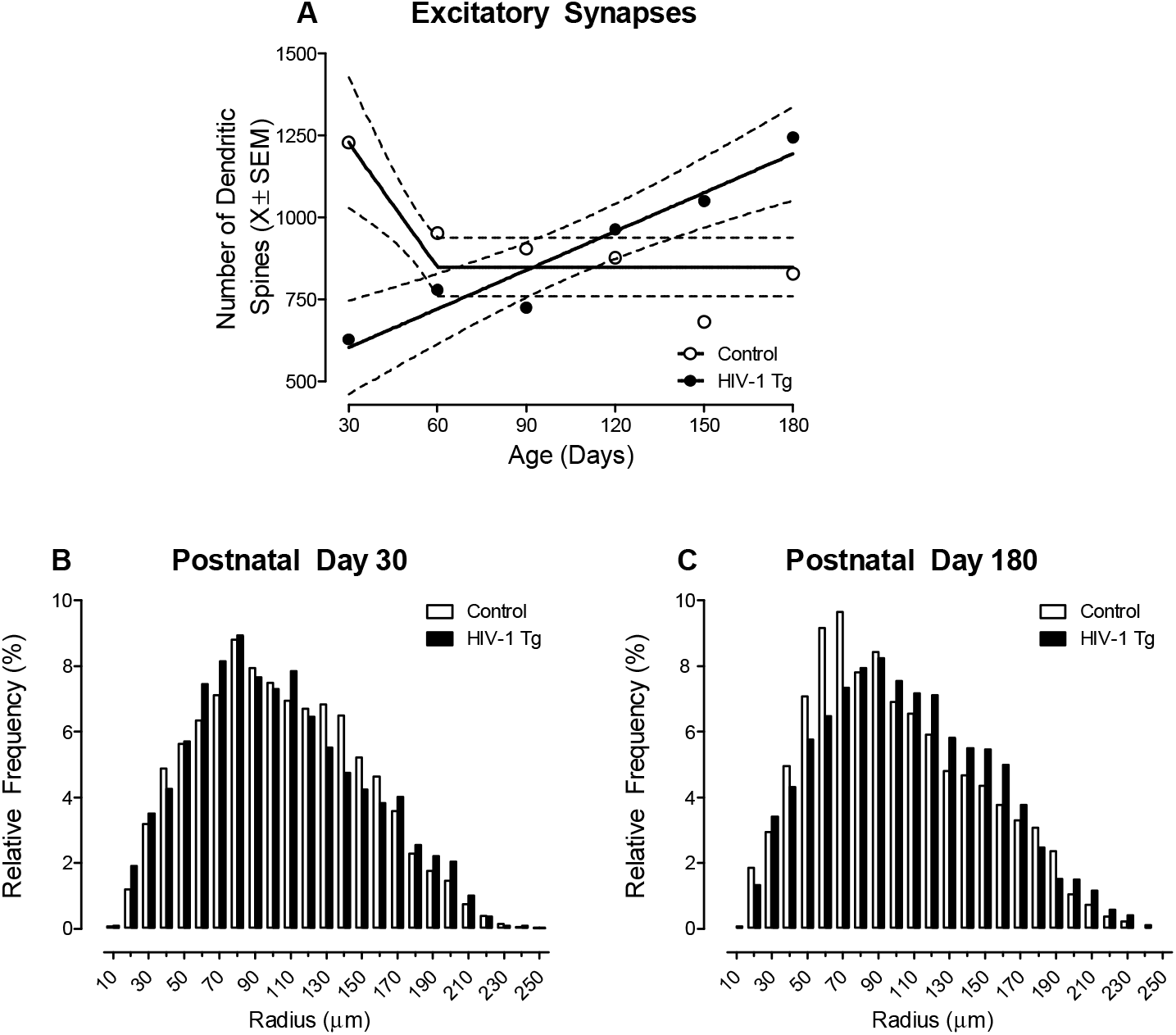
Prominent alterations in synaptogenesis and synaptic connectivity were observed in HIV-1 Tg rats. (**A**) The total number of dendritic spines (X ± 95% confidence intervals), a proxy for excitatory synapses [60], increased in a linear fashion in HIV-1 Tg animals. In sharp contrast, control animals exhibited an exuberant number of dendritic spines early in development (i.e., Postnatal Day (PD) 30); spines which were subsequently pruned during adolescence. (B,C) The distribution of the entire population of dendritic spines on pyramidal neurons from layers II-III of the medial prefrontal cortex are illustrated as a function of genotype (Control vs. HIV-1 Tg) and radii at PD 30 (**B**) and PD 180 (**C**). HIV-1 Tg animals displayed an increased relative frequency of dendritic spines on more distal dendrites relative to control animals at both PD 30 and PD 180; albeit the magnitude of the rightward shift increased with age, supporting a developmental decrease in synaptic connectivity.

#### 3.1.3 The magnitude of the rightward shift in the distribution of dendritic spines along the apical dendrite progresses with age in HIV-1 Tg animals, supporting a developmental decrease in synaptic connectivity

HIV-1 Tg animals exhibited a prominent rightward shift in the distribution of dendritic spines along the apical dendrite relative to control animals (Figure 3B-C; Genotype x Age x Radii Interaction, *F*(1,1816)=40.3, *p*≤0.001). Specifically, independent of age, HIV-1 Tg animals displayed a preponderance of dendritic spines on more distal dendrites relative to control animals; albeit a more pronounced rightward shift was observed at PD 180. Furthermore, it is notable that the factor of biological sex also influenced the magnitude, but not the pattern, of the distributional shift (Genotype x Age x Sex x Radii Interaction, *F*(1,1816)=154.6, *p*≤0.001). Results support, therefore, a developmental decrease in synaptic connectivity in HIV-1 Tg animals.

#### 3.1.4 HIV-1 Tg rats exhibited a progressive decrease in synaptic efficacy, indexed using dendritic spine volume

Strong correlations between dendritic spine volume and area of the postsynaptic density (PSD; [54; 61-62]) support a tight coupling between dendritic spine morphology and function. HIV-1 Tg animals exhibited a progressive decrease in dendritic spine volume throughout development; a sharp contrast to the age-related increase observed in control animals (Figure 4A; Best Fit: First-Order Polynomial, R^2^s≥0.54).

**Figure 4.**
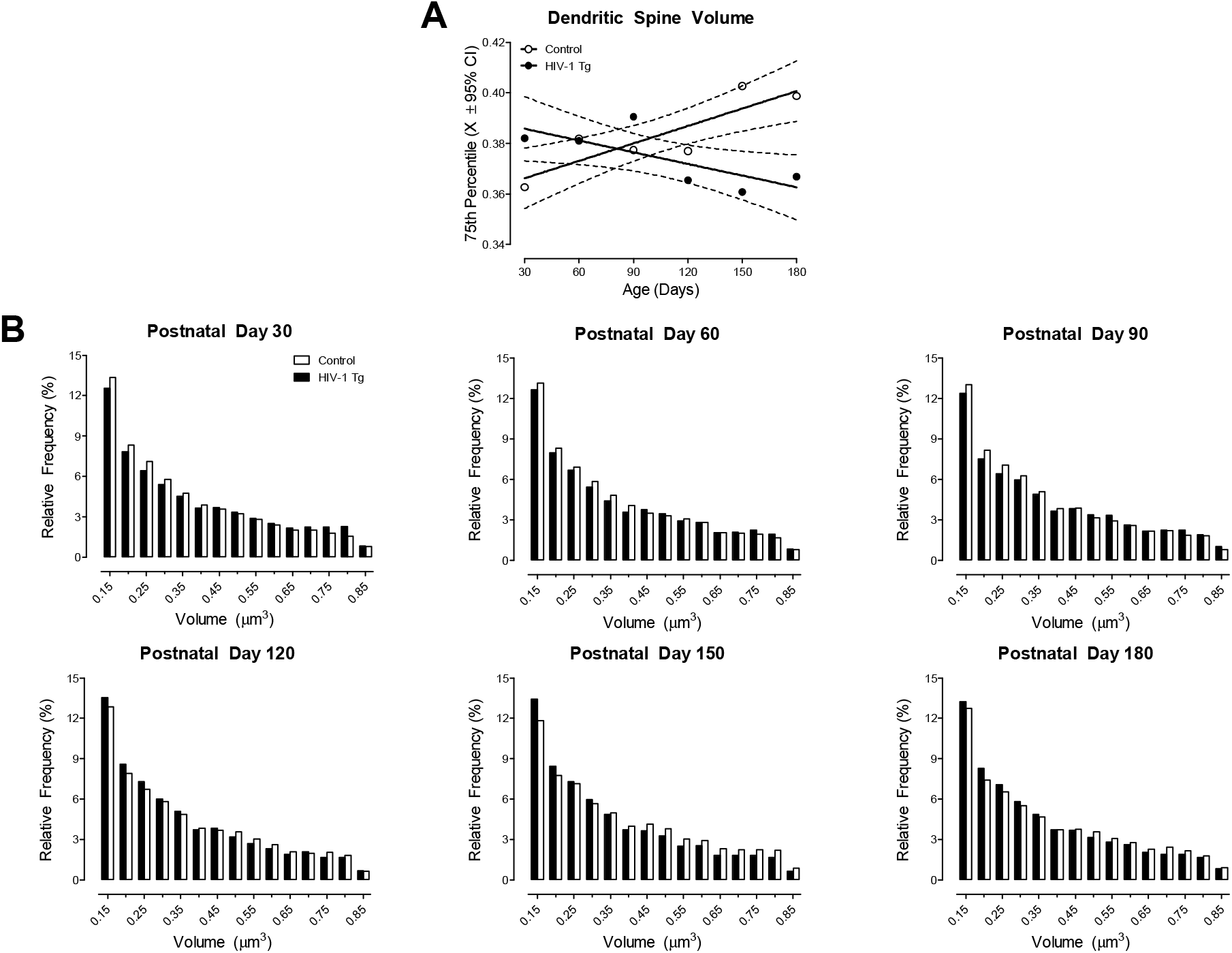
HIV-1 Tg animals exhibited a progressive decrease in synaptic efficacy, indexed using dendritic spine volume. (**A**) The 75^th^ percentile of dendritic spine volume was derived from the distribution of dendritic spines (X ± 95% confidence intervals). HIV-1 Tg animals exhibited a linear decrease in dendritic spine volume across development, whereas control animals displayed an age-related linear increase in dendritic spine volume. (**B**) Examination of the distribution of dendritic spine volume in HIV-1 Tg and control animals as a function of age confirmed the observations inferred from the 75^th^ percentile of dendritic spine volume.

Furthermore, examination of the distribution of dendritic spines (Figure 4B) confirmed the differential development of dendritic spine volume (Genotype x Age x Bin Interaction, *F*(80, 3200)=3.2, *p*≤0.001). Specifically, at PD 30, PD 60, and PD 90 HIV-1 Tg animals exhibited a prominent population shift towards increased dendritic spine volume relative to control animals. At PD 120, PD 150, and PD 180, however, a profound population shift towards decreased dendritic spine volume was observed in HIV-1 Tg, relative to control, animals. The factor of biological sex altered the magnitude (Genotype x Sex x Age x Bin Interaction, *F*(80,3200)=1.7, *p*≤0.001), but not the pattern, of the distributional shift. Collectively, HIV-1 Tg rats, independent of biological sex, displayed an age-related decrease in dendritic spine volume supporting a progressive loss in synaptic efficacy.

#### 3.1.5 Across time, the morphological parameters of dendritic spines in HIV-1 Tg animals regress, evidenced by a profound shift towards an immature phenotype

Additional dendritic spine morphological parameters, including dendritic spine length [63] and the dendritic spine head [54; 61-62], have also been associated with synaptic strength and area of the PSD, respectively. Examination of the distribution of dendritic spine backbone length (Figure 5A) and head diameter (Figure 5B) revealed a differential development of dendritic spine morphology (Genotype x Age x Bin Interaction, *F*(80, 3200)=14.8, *p*≤0.001 and *F*(55, 2200)=28.3, *p*≤0.001, respectively). Specifically, at PD 30, PD 60, and PD 90 HIV-1 Tg animals exhibited a prominent population shift towards shorter dendritic spines with increased head diameter relative to control animals. At PD 120, PD 150, and PD 180, however, a profound population shift towards longer dendritic spines with decreased head diameter observed in HIV-1 Tg, relative to control, animals. The predominant dendritic spine morphology in HIV-1 Tg animals, therefore, regresses from a mature ‘mushroom’ phenotype (i.e., large dendritic spine head and a small dendritic spine neck [64]) to an immature ‘thin’ phenotype (i.e., long, thin dendritic spine neck with a smaller dendritic spine head [64]) across time; a phenotype which contains smaller excitatory synapses [65].

**Figure 5.**
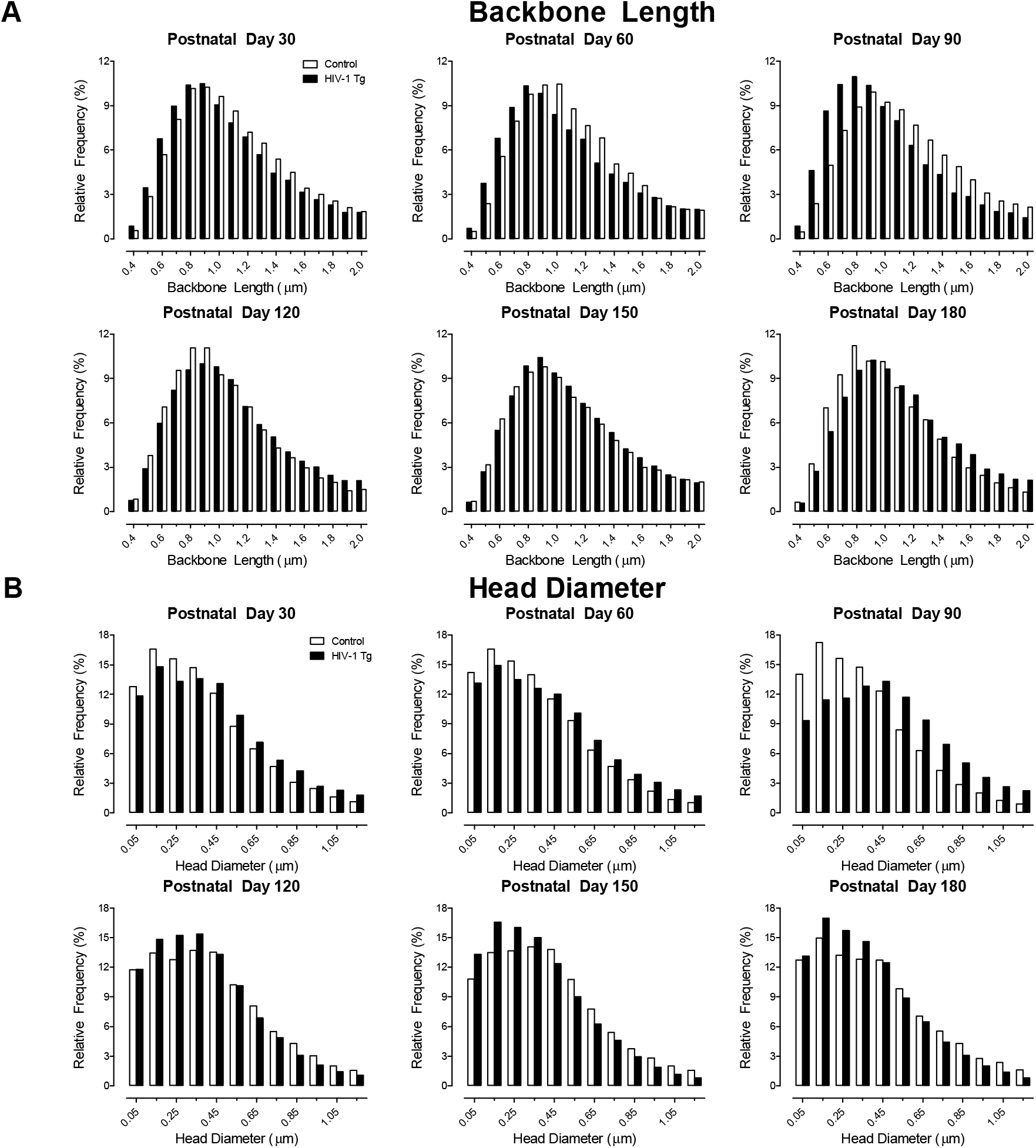
Dendritic spine morphology in pyramidal neurons from layers II-III of the medial prefrontal cortex in the HIV-1 Tg rat was further characterized by a developmental increase in dendritic spine backbone length (**A**) and progressive decrease in dendritic spine head diameter (**B**) relative to control animals. Collectively, HIV-1 Tg animals exhibited a morphological shift towards parameters consistent with a more immature dendritic spine phenotype across development. Data are illustrated as relative frequencies.

### 3.2. Experiment #2: Neuroinflammatory Markers in the Medial Prefrontal Cortex

#### 3.2.1 There is no compelling evidence for neuroinflammation in the mPFC of HIV-1 Tg rats during early development

Three putative neuroinflammatory markers, including IL-1β, IL-6, and TNF-α, were examined at PD 30 using RT-PCR. Overall, independent of genotype and/or biological sex, low levels of gene expression were observed, evidenced by cycle threshold (CT) values ranging from 26.1 to 40 for IL-1β, from 31.3 to 40 for IL-6 and from 25.7 to 40 for TNF-α. Data are presented as ΔCT values (i.e., Internal Control (β-actin) – Gene of Interest), whereby higher ΔCT values indicate lower gene expression and lower ΔCT values represent higher gene expression. With regards to IL-1β, a higher level of gene expression was observed in female animals, independent of genotype, relative to male animals (Figure 6A; Main Effect: Sex, *F*(1,35)=5.5, *p*≤0.03). Examination of IL-6 revealed a statistically significant main effect of genotype (Figure 6B; *F*(1,29)=6.2, *p*≤0.02), whereby HIV-1 Tg animals, independent of biological sex, exhibited a lower level of gene expression relative to control animals. For TNF-α gene expression was dependent upon an interaction between genotype and biological sex (Figure 6C; *F*(1, 33)=5.6 *p*≤0.02); an interaction resulting from decreased gene expression in HIV-1 Tg male animals relative to control male rats (*t*(33)=-2.2, *p*≤0.03).

**Figure 6.**
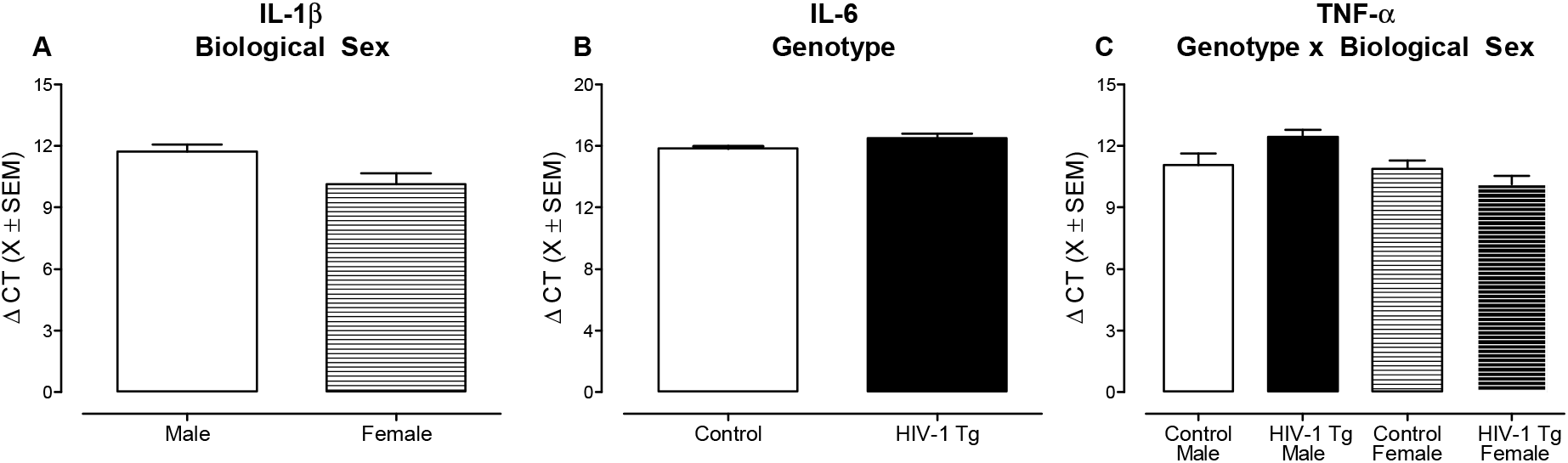
There was no compelling evidence for neuroinflammation in the medial prefrontal cortex of the HIV-1 Tg rat. Three putative neuroinflammatory markers, including IL-1β (**A**), IL-6 (**B**), and TNF-α (**C**) were assessed using real-time PCR (X ± 95% confidence intervals). Data are presented as ΔCT values (i.e., Internal Control (β-actin) – Gene of Interest), whereby higher ΔCT values indicate lower gene expression and lower ΔCT values represent higher gene expression. (**A**) With regards to IL-1β, a higher level of gene expression was observed in female animals, independent of genotype, relative to male animals. (**B**) A statistically significant main effect of genotype was observed for IL-6, indicating lower levels of gene expression in HIV-1 Tg animals relative to controls. (**C**) For TNF-α, a statistically significant genotype x sex interaction was observed; an interaction resulting from decreased gene expression in HIV-1 Tg male animals relative to control male rats.

Given that the highest levels of gene expression were observed for IL-1β, ELISA was subsequently performed to evaluate IL-1β at the protein level. There were no statistically significant main effects and/or interactions for the concentration of IL-1β (*p*>0.05; Supplementary Figure 3). Thus, collectively, there is no compelling evidence for neuroinflammation in the mPFC of HIV-1 Tg rats.

## 4. Discussion

Constitutive expression of HIV-1 viral proteins induces prominent neurodevelopmental alterations, independent of biological sex, in pyramidal neurons from layers II-III of the mPFC. Profound alterations in dendritic and synaptic pruning, regressive processes that are a hallmark of adolescent brain maturation [for review, 66], were observed in HIV-1 Tg rats. A developmental decrease in synaptic connectivity was evidenced by an increased prevalence of dendritic spines on more distal branches of the apical dendrite with age. Morphological parameters of dendritic spines further revealed an age-related progressive decrease in synaptic efficacy characterized by a developmental shift towards an immature dendritic spine phenotype. Based on an examination of three putative neuroinflammatory markers early in development (i.e., PD 30), there was no compelling evidence for neuroinflammation in the mPFC of HIV-1 Tg rats. Understanding the neural mechanisms underlying chronic neurocognitive impairments in pALHIV may afford key targets for innovative therapeutics and cure strategies; an urgent need given the growing population of pALHIV.

In the PFC, neurite [17; 67] and synaptic [15; 22] pruning are a hallmark of adolescent neurodevelopment [66]. Observations in control animals, independent of biological sex, corroborates these previous findings. In HIV-1 Tg animals, however, regressive processes were profoundly altered, evidenced by a linear increase in indices of neuronal arbor complexity, dendritic branching complexity, and excitatory synapses throughout development; alterations which may be due, at least in part, to microglial dysfunction. Specifically, early in the course of infection, HIV-1-infected monocytes migrate across the blood-brain barrier infiltrating the brain and infecting microglia [68-69]. Following chronic HIV-1 viral protein exposure, microglia serve as a viral reservoir for HIV-1 in the brain [69-71] supporting a cell type that may underlie aberrant neurite and synaptic pruning.

Under homeostatic conditions, microglia exhibit a ramified morphology with small somas and highly branched processes supporting constant environmental surveillance [72-73]; surveillance which is uniquely targeted to synaptic structures, including dendritic spines [74-75]. During brain development, microglia are involved in the phagocytic and/or trogocytotic elimination of neurites [e.g., 76-78] and synapses [e.g., 79-81]. Proteins and receptors expressed by microglia (e.g., complement component 1q (C1q) [82]; complement receptor 3 (CR3) [83]; CX3C chemokine receptor 1 (*Cx3cr1*) [84-85]), have been implicated as potential mechanisms underlying microglia-mediated neurite and pre- and postsynaptic engulfment [77-80; 86]. Specifically, genetic deletion of C1q [86], CR3 [80], or *Cx3cr1* [77; 79] precludes neurite and/or synaptic pruning. Furthermore, increasing or decreasing the activity level of the complement system results in either excessive or impaired neurite pruning, respectively [78]. Critically, the complement system and *Cx3cr1* are dysregulated by HIV-1 viral proteins [87-88] and associated with neuronal injury in HIV-1 seropositive individuals [89]; alterations which support a convergent mechanism underlying the prominent alterations in regressive processes observed in the HIV-1 Tg rat.

After synaptic pruning, the remaining synapses are functionally strengthened and elaborated to aid in the formation of mature neural circuits. The PFC is highly connected with other cortical and subcortical regions making it well suited to serve as a central hub, integrating and relaying information throughout the brain [for review, 90]. In addition to excitatory glutamatergic and inhibitory gamma-aminobutyric acid afferents, the PFC is densely innervated by dopaminergic, noradrenergic, and serotonergic projections from multiple brain regions (i.e., ventral tegmental area, locus coeruleus, and dorsal and median raphe nuclei, respectively [18]). The vast majority of afferent projections synapse on dendritic spines of pyramidal neurons [91]. Therefore, two complementary approaches were utilized to establish progressive neural circuitry dysfunction in the HIV-1 Tg rat, including an examination of the distribution of dendritic spines and evaluation of dendritic spine morphological parameters.

Independent of age (i.e., PD 30 or PD 180), HIV-1 Tg animals exhibited a prominent alteration in synaptic connectivity, evidenced by a prominent rightward shift in the distribution of dendritic spines along the apical dendrite relative to controls. Architecturally, the PFC is organized in a laminar fashion from superficial (i.e., Layer I) to deep (i.e., Layer VI [18]); pyramidal neurons from layers II-III of the mPFC extend superficially into layer I. Dopaminergic, noradrenergic, and serotonergic receptors are abundantly expressed in layers II-III, but not layer I, of the mPFC [92]. In HIV-1 Tg rats, therefore, the preponderance of dendritic spines on distal dendrites, which extend into layer I, supports alterations in neurotransmitter innervation and synaptic connectivity. Indeed, neurotransmitter systems, including the dopamine [e.g., 93-96], norepinephrine [e.g., 97] and serotonin [e.g., 98-99] systems, are dysregulated by HIV-1 viral protein exposure. Furthermore, the magnitude of the rightward shift in the distribution of dendritic spines along the apical dendrite progresses with age in HIV-1 Tg animals, supporting an age-related loss in neurotransmitter innervation and synaptic connectivity.

Examination of dendritic spine morphological parameters adds additional support for the progressive decrease in neurotransmitter innervation and synaptic connectivity. A discrete classification system, consisting of four dendritic spines categories (i.e., filopodia, mushroom, stubby, thin), has been classically utilized to characterize the morphology of dendritic spines [64]. More recently, however, it has been recognized that spine morphology is more accurately assessed by continuous numerical measures [e.g., 54-55; 100]. Indeed, the utilization of continuous morphological measurements has revealed the tight coupling between spine structure and synaptic function, whereby total spine volume and dendritic spine heads are positively correlated with the area of the PSD [54; 62] and neck length is negatively correlated with synaptic efficacy [63]. Critically, a pronounced PSD, located at the tip of dendritic spines, is a primary characteristic of excitatory glutamatergic synapses [for review, 101]. Indeed, the size and glutamate sensitivity of the PSD is correlated with AMPA receptor immunoreactivity [102-103] and the dendritic spine head [104], respectively. HIV-1 Tg animals exhibited an age-related decrease in dendritic spine volume and head diameter, as well as an age-related increase in backbone length; a population shift supporting a developmental decrease in the area of the PSD and synaptic efficacy. Collectively, the investigation of dendritic spines supports the dysregulation of both monoaminergic and glutamatergic neurotransmission.

Notably, the neurodevelopmental alterations in regressive processes and synaptic function occurred in the absence of any significant neuroinflammation early in development; findings which are consistent with our current knowledge of the clinical syndrome in the post-cART era. Inflammatory markers measured in the periphery (e.g., blood samples) of children perinatally infected with HIV-1 are significantly elevated relative to controls [e.g., 105-107]. Peripheral inflammation, however, is distinct from neuroinflammation (e.g., measurements from cerebrospinal fluid (CSF)) evidenced by limited concordance between plasma and CSF immunological markers in children perinatally infected with HIV-1 [107]. Indeed, in the CSF of HIV-1 seropositive children, multiple proinflammatory markers are undetectable [108]. We recognize that the proposed role of microglia in regressive processes and synaptic dysfunction may seem contradictory given the relative absence of neuroinflammation. Therefore, we hypothesize that chronic HIV-1 viral protein exposure results in microglial dysfunction, evidenced by cellular senescence [109].

The persistence of progressive chronic neurocognitive impairments necessitates the development of efficacious adjunctive therapeutics; inferences drawn from the present study have revealed three potential therapeutic targets. First, despite the great promise of complement-targeted therapeutics for neurological diseases, the safety of targeting the complement pathway is of utmost concern given its importance in host immune defenses [for review, 110]. Second, the therapeutic efficacy of targeting monoaminergic [e.g., 111-113] or glutamatergic [e.g., 114-116] neurotransmission for chronic neurocognitive impairments and/or HAND has been inconsistent. Third, therapeutics targeting chemokine receptors are limited by small sample sizes (*n*<20 [117-118]), primarily male samples [117-119], and inconsistent improvements in cognitive function [117-119]. Fourth, therapeutics that exert their effects by enhancing synaptic function, have, at least in preclinical studies, shown greater promise for chronic neurocognitive impairments [120] and HAND [113; 115; 121-124].

In conclusion, constitutive expression of HIV-1 viral proteins induced profound neurodevelopmental alterations, independent of biological sex, in pyramidal neurons from layers II-III of the mPFC. Neurodevelopmental alterations in the HIV-1 Tg rat were characterized by aberrant neurite and synaptic pruning, a progressive decrease in synaptic connectivity, and an age-related population shift towards a more immature dendritic spine phenotype. Understanding the neural mechanisms underlying chronic neurocognitive impairments in pALHIV may afford a key target for innovative therapeutics and cure strategies; an urgent need given the growing population of pALHIV.

## Supplementary Materials

The following are available online, Figure S1: Body Weight, Figure S2: Sholl Intersection Profile, Figure S3: IL-1β ELISA.

## Author Contributions

R.M.B and C.F.M. designed the study. K.A.M., H.L., R.M.B., and C.F.M. planned the study and collected the data. K.A.M., H.L., and C.F.M. analyzed the data. K.A.M., R.M.B., and C.F.M. wrote the manuscript. All authors critically appraised and approved the final version of the manuscript.

## Institutional Review Board Statement

Not applicable.

## Data Availability Statement

All relevant data are within the manuscript.

