## Supplementary Figures 1-3 for "Neurodevelopmental processes in the prefrontal cortex derailed by chronic HIV-1 viral protein exposure"

Program in Behavioral Neuroscience

Department of Psychology

University of South Carolina

Columbia, SC 29208

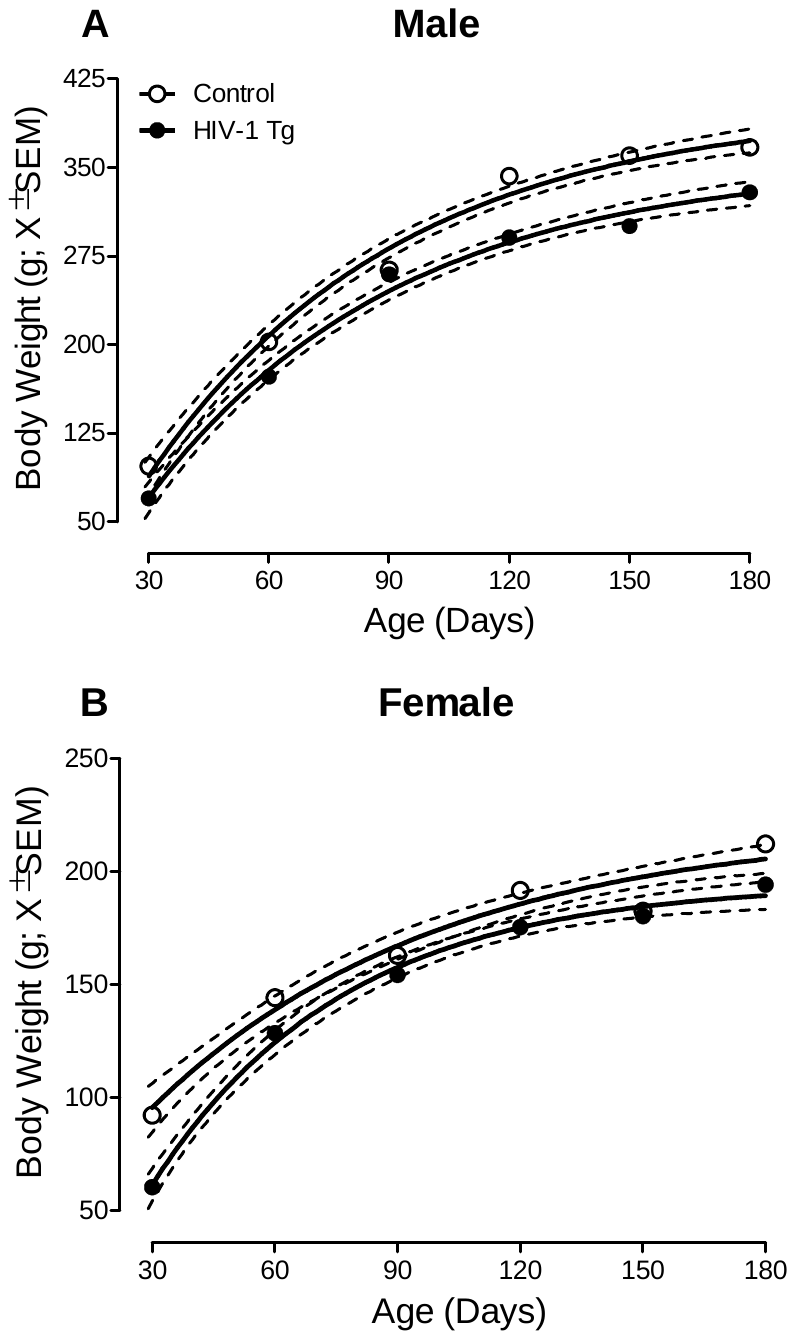

**Figure S1: Body Weight.** Mean body weight, a measure of somatic growth, is illustrated for both males (**A**) and females (**B**) as a function of genotype (Control vs. HIV-1 Tg) and age (X ± 95% confidence intervals). Independent of biological sex, HIV-1 Tg animals weighed significantly less than their control counterparts; no statistically significant differences in the rate of growth were observed.

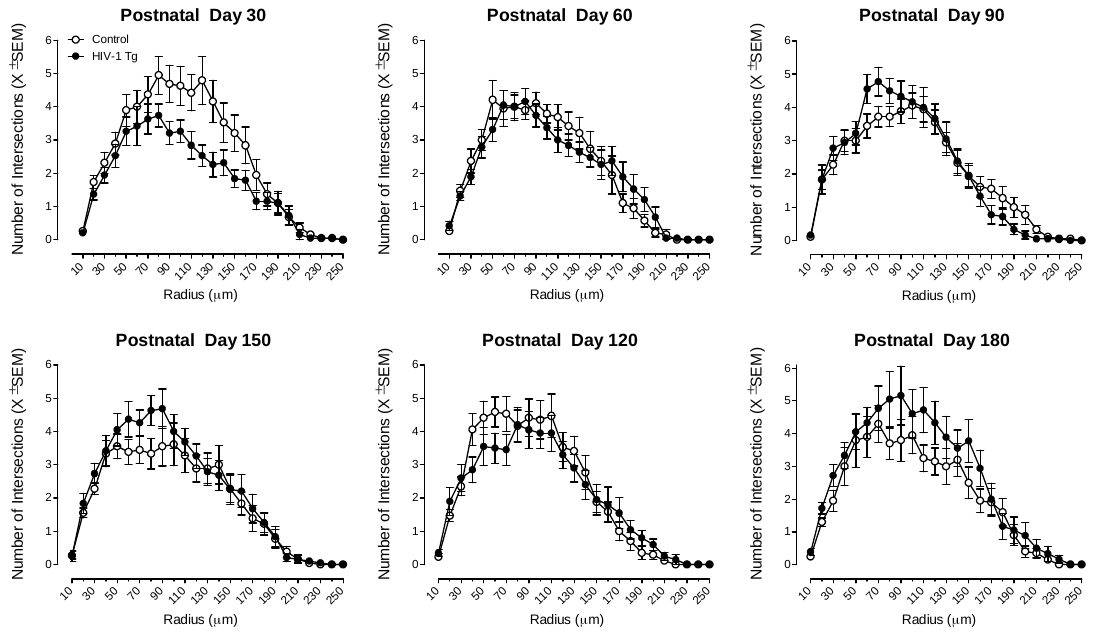

**Figure S2: Sholl Intersection Profile.** The classic Sholl intersection profile is illustrated as a function of genotype (Control vs. HIV-1 Tg) and age (X ± SEM). At postnatal day (PD) 30, control animals exhibited exuberant neuronal arbor complexity in pyramidal neurons from layers II-III of the medial prefrontal cortex relative to HIV-1 Tg animals. A reduction in neuronal arbor complexity was observed in control animals at PD 60 consistent with adolescent dendritic pruning. In sharp contrast, neuronal arbor complexity increased in HIV-1 Tg animals throughout development. Results support, therefore, aberrant neurite pruning in HIV-1 Tg animals.

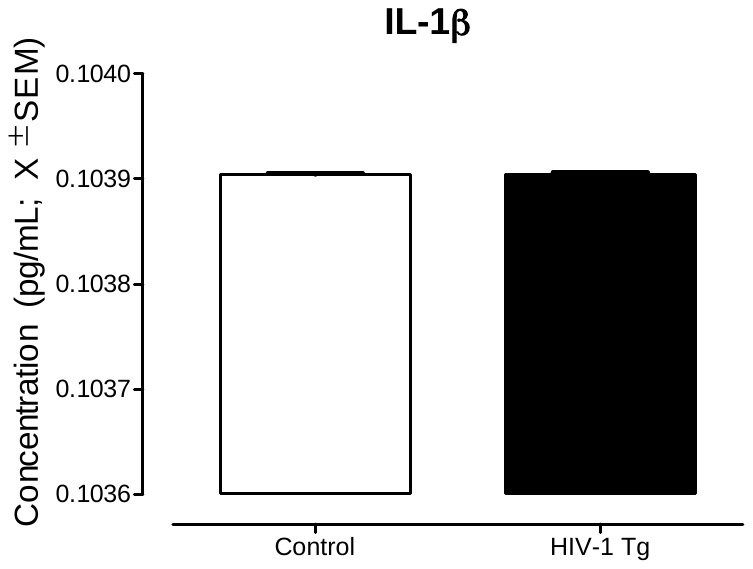

**Figure S3: IL-1β ELISA.** ELISA was conducted in the prefrontal cortex at postnatal day 30 to evaluate 1L-1β protein levels. No statistically significant main effects and/or interactions were observed for the concentration of IL-1β. Data are presented as concentration (pg/mL; X ± SEM) dependent upon genotype (Control vs. HIV-1 Tg).
